# “Eating to Survive… or not? feeding compensation and toxicity in a songbird exposed to thiamethoxam treated seeds”

**DOI:** 10.1101/2025.11.25.690477

**Authors:** Maria Belen Poliserpi, Elena Fernandez-Vizcaino, Julie Celine Brodeur

**Affiliations:** Instituto de Recursos Biológicos, Centro de Investigaciones de Recursos Naturales (CIRN). Instituto Nacional de Tecnología Agropecuaria (INTA), (1686) Hurlingham, Buenos Aires, Argentina; Departamento de Zoología, Facultad de Ciencias, Universidad de Córdoba, Córdoba, Spain; Instituto de Investigación en Recursos Cinegéticos (CSIC-UCLM-JCCM), Departamento de Toxicología, Ciudad Real, Spain; Consejo Nacional de Investigaciones Científicas y Técnicas (CONICET), Argentina

**Keywords:** Pesticides, Neonicotinoids, Seed treatment, Bird, Behavior, Agriculture

## Abstract

Thiamethoxam (TMX) is a neonicotinoid insecticide widely used for seed treatment in agriculture. Birds can be exposed to TMX by ingesting unburied seeds after sowing. This study evaluated the toxicological effects of such an exposure scenario in a common farmland passerine *Agelaioides badius*. Birds were subjected to a 21-day dietary exposure of TMX-treated seeds at 0 (control), 0.027 (low), 0.33 (medium), and 4.3 (high) g TMX/kg seed (N=8 per group), representative of typical agricultural practices. A dose-dependent increase in seed consumption was observed in the latter stages of the experiment, suggesting compensatory feeding that amplifies toxicant intake. Notably, the high-dose group exhibited 50% mortality, an unexpected finding of concern given TMX’s classification as of low acute toxicity. High-dose birds displayed significant body mass reduction (−20.5%) and impaired anti-predator responsiveness. Furthermore, general activity patterns were altered across all treated groups. At a physiological level, TMX exposure in the high-dose group induced reduced hematocrit and decreased polychromasia. Tissue-specific modulations in enzymatic biomarkers, including elevated cholinesterase and glutathione S-transferase activities, indicated oxidative stress and cholinergic alterations. These findings demonstrate that environmentally-relevant ingestion of TMX-coated seeds can elicit sublethal and lethal effects in granivorous birds, potentially posing a significant risk to farmland bird populations.

Thiamethoxam effects on birds are little studied due to its reported low acute toxicity. This study demonstrates that ingestion of thiamethoxam-treated seeds causes multilevel biological effects and compensatory feeding, underscoring ecological risk for farmland bird populations.

## INTRODUCTION

Seed treatment has become an essential part of modern agriculture (Hahne et al., 2024). Neonicotinoids are a class of insecticides widely used in seed treatment due to their systemic action that enables protection of the entire plant, and their greater toxicity to insects compared to vertebrates (Tomizawa and Casida, 2005). Since the adoption of neonicotinoids, the evidence of their negative effects on pollinators and other species (Main et al., 2021; Molenaar et al., 2024), together with their widespread presence in the environment (Pietrzak et al., 2020) led to limitations in their use. While some regions like Europe (Regulation 2018/783–785) and Canada restricted their use (Health Canada, 2019), the treatment of seeds with neonicotinoids is still practiced worldwide (Pearsons et al., 2021; Thompson et al., 2020).

The ingestion of coated crop seeds represents the principal route of exposure to neonicotinoids in wild birds, at levels that can cause adverse effects and mortality (Brodeur and Poliserpi, 2023; Poliserpi et al., 2021b). Globally, the most used neonicotinoids are imidacloprid (IMI), thiamethoxam (TMX), and clothianidin (CLO) (Bass et al., 2015; Jeschke et al., 2011). To birds, IMI can be over 70X times more toxic than TMX or CLO (Addy-Orduna et al., 2019). For this reason, the effects of IMI have been more extensively studied in comparison to that of other neonicotinoids (Mineau, 2023). The toxicity of neonicotinoids is linked to their interaction with the nervous system, where they act as nicotinic acetylcholine receptor (nAChR) agonist (Tomizawa and Casida, 2005), (Gibbons et al., 2015; Lopez-Antia et al., 2013; Poliserpi and Brodeur, 2023). Beyond mortality, a range of sublethal effects were documented in birds after exposure to neonicotinoids, including behavioral, physiological, and reproductive (Gibbons et al., 2015; Mineau and Palmer, 2013; Pisa et al., 2017).

Neonicotinoid exposure in birds is frequently associated with body weight loss, often linked to reduced food intake and appetite suppression, as observed with IMI, and less clear for TMX and CLO (Addy-Orduna et al., 2022; Eng et al., 2019; Eng & Morrissey, 2025). Beyond appetite suppression, neurotoxic effects may disrupt feeding behavior, increasing feeding time without greater consumption, as suggested for songbirds fed with IMI-treated seeds (Brodeur and Poliserpi, 2023; Poliserpi et al., 2023).

The disruption of essential behaviors such as reproductive, anti-predatory, and migration was observed in birds exposed to neonicotinoids, particularly to IMI (Addy-Orduna et al., 2024; Eng et al., 2019; Poliserpi and Brodeur, 2023). Behavioral impairments are considered sensitive indicators of contaminant exposure and often precede severe physiological effects (Ågerstrand et al., 2020; Ford et al., 2021; Grunst et al., 2023). In birds and mammals, some behavioral changes related to movement were linked to cholinergic dysfunction after exposure to neonicotinoids (Abu Zeid et al., 2019; Kapoor et al., 2014; Rodrigues et al., 2010); while behaviors related to high energy demand were associated with oxidative stress and alteration in hematological profiles (Bal et al., 2012; Jenni-Eiermann et al., 2014; Lopez-Antia et al., 2013; Pap et al., 2018; Wang et al., 2018).

The neonicotinoid thiamethoxam is classified as moderately to slightly hazardous (WHO class II and III), and is considered of low acute, short-term and long-term toxicity to birds (FAO, 2021). However, is of particular interest because it is metabolized into CLO another neonicotinoid, and both compounds have been shown to cause adverse effects in birds (Fernández-Vizcaíno et al., 2025; Pan et al., 2022a). The occurrence of TMX residues in wild birds highlights the need for further research on its effects to refine risk assessment (Humann-Guilleminot et al., 2019; Klaas-Fábregas et al., 2024; Nimako et al., 2025). The grayish baywing (*Agelaioides badius*) has been previously validated as a suitable model species for avian ecotoxicology (Brodeur and Poliserpi, 2017; Poliserpi and Brodeur, 2023). This granivorous passerine, classified as “Least Concern” (BirdLife International, 2021) is abundant in agricultural landscapes of the Americas and has a diet composed predominantly of seeds (Frutos et al., 2016) making it particularly vulnerable to neonicotinoids-treated seeds. In this context, the present study aimed to evaluate the effects of dietary exposure to TMX-treated seeds in *A. badius* over a 21-day period, using concentrations representative of those applied in the Pampa region of Argentina (i.e., 0.03-3 g IMI/kg seed). The following aspects were examined: 1) quantify the weekly consumption of TMX-treated seeds offered *ad libitum*, 2) assess behavioral alterations associated with TMX ingestion, and 3) evaluate hematological and biochemical responses indicative of physiological stress and toxicity exerted by TMX.

## 2. METHODOLOGY

### 2.1. Capture and housing conditions

Wild adult grayish baywings older than a year (N=32)(Fraga, 1991) were captured within the limits of the National Center for Agricultural Research (CNIA), located in Hurlingham, Buenos Aires Province, Argentina (34◦36′24″ S, 58◦39′56″ W). Detailed information on the biology, capture and husbandry of grayish baywing is provided by (Brodeur and Poliserpi, 2017). Baywings were captured with baited funnel traps during the non-reproductive season (i.e., May to August, the fall-winter season in the South Hemisphere) with the proper permits (Flora and Fauna Department of Buenos Aires Province, Argentina (Resolution #22500–38,407/17-19).

The birds were acclimatized to laboratory conditions for 15 days: temperature was maintained at 23 ± 2°C and the photoperiod was set at 10:14 h light: dark. Birds were provided unchlorinated well water and fed with a commercial seed mixture containing birdseed, rapeseed, chia, oat, millet and flax seed enriched with calcium, phosphorus, and vitamins A, B1, B2, B6, B12 (food composition: protein 17,4 %, fat 8 %, fiber 9.5 %, ashes 3.5 %, humidity 12 %; commercialized by Ibis, Nor S.A, Argentina). After the acclimatation period, the food was changed to peeled millet (Forrajera Lunic, Argentina) for 5 days before conducting the experiment. Peeled millet was selected as experimental seed to avoid dehusking, and to ensure a precise determination of ingested doses.

### 2.2. Seed preparation and exposure concentrations

Peeled millet seeds were treated with the commercial formulation Actara^®^ (Syngenta). The formulate consists of a water dispersible granule containing 25% of TMX 3-(2-cloro-tiazol-5-ilmetil)-5-metil- [1,3,5] oxadiazinan-4-ilideno-N-nitroamina; CAS No. 153719–23-4. The formulation contains proprietary surfactants, additives or emulsifiers of undisclosed molecular nature.

Three experimental treatments and a control were defined to insure that long-term sublethal effects could be examined. The concentrations selected were 0 (Control), 0.03 (Low), 0.3 (Medium), and 3 (High)g TMX/kg seed, which cover the range of concentrations approved for seed coating with TMX in Argentina (Table S1, SENASA, 2024). TMX solutions were prepared by diluting the formulate in distilled water following the instructions provided by the manufacturer. The experimental seeds were prepared by spraying over the peeled millet while continuously mixing the seeds in a flat container, to ensure a uniform coating. Sprayed millet was left to air dry at room temperature overnight then divided into 10 g aliquots and stored at −20 °C until used. Experimental seeds were prepared weekly to avoid degradation of the active ingredient. A total of three batches of seeds were prepared for each experimental group.

Analytical verification of TMX was determined as described in (Poliserpi et al., 2023). Briefly, 1 g of seeds was extracted by salting out with 0.5 g of sodium acetate, and 2 g of magnesium sulfate, together with 1 mL water, and 10 mL acetonitrile containing acetic acid (1 %). The mixture was placed in an ultrasonic bath for 30 min and centrifuged 10 minutes at 3500 rpm. One mL of the supernatant was diluted with 1 mL of HPLC-grade water and filtered through a 0.22 μm nylon membrane. Thiamethoxam PESTANAL® (Merck, Alemania) analytical standard was used for calibration curves and quality control samples by fortifying control seeds. TMX determination was carried out with an Acquity UPLC BEH C18 column (1.7 μm, 100 × 2.1 mm). Detection and quantification limits were 0.006 and 0.01 μg/g, respectively, and the standard curve presented strong linearity (r^2^ = 0.998). Average TMX concentration per treatment (N=3) was (mean ± S.E.): low = 0.027 ± 0.0033; medium= 0.334 ± 0.0334, and high= 4.34 ± 0.667 g TMX/kg seeds. No TMX was detected in control seeds.

### 2.3. Experimental design

The birds were fed for 21 days with control or TMX-treated seeds *ad libitum*. Because baywings are monomorphic, sex was determined after the experiment. Eight birds were assigned to each experimental group. Every morning between 9:00 and 11:00 a.m., 10 g of treated seeds were placed into the food container of each bird, and on the next morning, the remaining seeds were collected and weighed to the nearest 0.001 g with an electronic balance. Seed intake was expressed weekly as g of seeds per kg of body weight (BW). The TMX daily dose ingested by each bird was calculated estimated with the average analytical concentration of TMX in the seeds and expressed as mg of TMX per kg of BW. Birds were weighted using a Pesola® spring scale (± 1 g) the day before the exposure and then once a week. Feeding behavior, activity and anti-predatory behavior, were evaluated by filming the birds once a week as detailed below. Survival of birds was assessed daily. At the end of the exposure, birds were mandatory euthanized in a carbon dioxide gas chamber in agreement with institutional and internationally accepted animal welfare guidelines (Fair et al., 2010). All procedures and protocols involving the birds were approved by the Institutional Animal Care and Use Committee (CICUAE-INTA permit #24/2021).

### 2.4. Feeding behavior and activity in the cage

Baywings were filmed undisturbed two times a day: the first and last hour of light (i.e., daytime: morning and afternoon), coinciding with the period of higher activity in birds (Sutherland et al., 2004).With the camaras (Dahua®) permanently placed in the laboratory, one-minute videos, spaced by five minutes, were obtained to ensure the independence of the registered events. Each day, twelve videos were selected (i.e., six for each daytime) per bird, and analyzed using ANY-maze Behavioral Tracking Software (Stoelting Co.). A bird was considered feeding when it was either with the head inside the food container or manipulating the seeds with the beak. To evaluate feeding behavior, the number of visits and the total time on the feeder was registered. Also, the average duration of the feeding events was calculated by dividing the total time in the feeder by the number of visits. The activity was measured as the movement of birds between the sectors of the cage: floor, perch and feeder: each transition was counted as a jump. The number of jumps per individual were registered.

### 2.5. Anty-predatory behavior

The response of the grayish baywings to predator vocalization was evaluated following (Poliserpi and Brodeur, 2023). Briefly, the vocalization of the Harris’s Hawk (*Parabuteo unicinctus*), a natural predator, was played in the laboratory while birds were undisturbed, and were video recorded (xeno-canto/49468). A score of four categories was defined based on the intensity of the anti-predatory response of a bird: (0) does not show any sign of listening the call; (1) listen to the call by slightly moving the head but does not alter their behavior; (2) stays vigilant in the place, scan its surroundings, stretch the neck or crouches; (3) the bird jumps, flies or makes alert calls. The response was also classified as whether the individuals reacted or not to the predator vocalization: “reaction” was considered when birds were scored with 2 or 3 (i.e., birds modify their behavior after the stimulus), and “no reaction” when birds had scores of 0 and 1 (i.e., birds did not show anti-predatory behavior).

### 2.6. Sample collection

Sample collection was conducted after mandatory euthanasia, following animal welfare guidelines, as previously described by Poliserpi and Brodeur (2023). Briefly, blood was obtained by cardiac punction in a heparinized syringe. A blood smear was prepared per individual for polychromasia (PC), and total white blood cell (WBC) counts. To estimate packed cell volume (PCV), blood was collected in two heparinized capillary tubes that were centrifuged at 10 000 g for 5 min. The remaining blood was centrifuged at 10 000 g for 5 min, and the plasma was separated and stored at −80°C. Multiple tissues were collected to conduct biochemical analysis. The liver was weighed to the nearest 0.001 g to calculate the hepatosomatic index (HSI) as the ratio of liver weight to total body weight (BW) at the end of exposure. Brain, liver, kidney, and pectoralis muscle samples, were collected and stored at −80°C until the analyses were performed. The sex of the birds was determined by visual examination of the gonads.

### 2.7. Hematological parameters and enzymatic activity

PCV was estimated as the percentage of the solid fraction of blood from the total volume. Blood smears were air-dried, fixed with methanol, and stained with Wright-Giemsa solution (Biopack^®^ commercial solution). Total WBC and PC were estimated per 1000 erythrocytes by counting the total number of WBC or polychromatophilic (immature) erythrocytes, respectively, in a monocellular layer examined at 1000X, among a total of 10.000 erythrocytes per individual (Campbell, 2015). Enzymatic activities of glutathione-S-transferase (GST), cholinesterase (ChE), and catalase (CAT) were determined in liver, muscle, brain, kidney, and plasma as previously described in (Poliserpi et al., 2021a; Poliserpi and Brodeur, 2023). Briefly, tissue samples were homogenized in ice-cold 50 mM tris (hydroxymethyl)aminomethane buffer (pH 7.4) containing 1 mM ethylenediaminetetraacetic acid and 0.25 M of sucrose. Homogenates were centrifuged at 10.000 g for 15 min at 4° C to remove debris, and the supernatant was used for enzymatic determinations. Plasma was directly diluted in PBS for enzymatic determinations.

### 2.8. Data analysis

R Software 4.5.1 was used for all statistical analysis, with exception of survival, for which GraphPad Prism 5.03 software was used (GraphPad Software, San Diego, California USA). Statistical significance was considered at the p<0.05 level, with p-values between 0.05 and 0.1 being considered as marginally significant. Continuous variables were checked for normality using with Kolmogorov-Smirnov tests. Those variables that did not follow normal distribution were transformed using the decimal logarithm (ChE Brain, GST Brain, GST Muscle, CAT kidney, ChE liver and CAT liver). The survival of baywings was analyzed using Mantel-Cox test for comparisons among treatment groups. To explore factors influencing weight and weekly consumption of seeds (grams of seeds/ kg BW), Generalized Linear Mixed Models (GLMMs) were employed. Treatment, week (3 levels), sex, and their interactions served as explanatory variables, with individuals included as a random factor. Within treatments, comparisons were made versus pre-exposure values (i.e. week 0). Anti-predatory behavior was evaluated with GLMMs with a binomial (i.e., predation reaction) or multinomial (i.e., predation reaction score) distribution and logit ordinal link function. Treatment, day, and their interaction served as explanatory variables, with individuals included as a random factor. To assess the treatment effect on behavioral parameters: number of feeding visits and jumping occurrences (discrete continuous variables), a GLMM with a Poisson distribution and a logarithmic link function was conducted. The behavioral variables: total time on the feeder, and duration of feeding events (continuous variables), GLMMs with normal distribution were used. Treatment, day, and daytime (morning or afternoon) were used as explanatory variables, with daytime nested within day, and individual as random effect. Factors affecting physiological parameters measured at the end of the experiment, like enzymatic activity and hematology parameters, were analyzed with general linear models (GLM), with the treatment and sex as fixed factor. A stepwise backward selection was used to eliminate model terms that did not significantly explain variations in the response variable, either alone or in interaction with other terms. The treatment factor was not considered for elimination from the models in order to report treatment effects. Birds that died during the exposure were not included in the analysis.

## 3. RESULTS AND DISCUSION

The experimental design of this study simulated environmentally relevant conditions by using TMX concentrations within the approved range for seed coating and an exposure period reflecting the availability of freshly treated seeds during the sowing season. This approach represents a realistic exposure scenario for birds, since the low and medium concentrations could correspond to situations where approximately 0.7% or 7% of the baywing’s diet consists of seeds coated with 4.3 g TMX/kg seed (i.e., the high-concentration treatment). Under this realistic exposure scenario, grayish baywings exhibited significant adverse effects at multiple levels: mortality occurred in the group exposed to the highest TMX concentration, surviving birds lost body weight, and all TMX-exposed individuals showed behavioral and biochemical alterations, as well as a dose-dependent increase in food intake.

### 3.1 TMX doses and survival

Thirty-two grayish baywings were used in the experiment: 17 females and 15 males with initial BW (mean ± S.D.) of 40.73 ± 2.64 g and no significant differences were observed sexes (sexes F (1) =2.127, p = 0.158; data not shown). Daily doses ingested by the birds (mean ± S.E.) were 2.961 ± 0.071, 38.61 ± 1.26 and 628.4 ± 14.03 mg TMX/kg BW for low, medium and high treatments, respectively. The survival of the birds from control, low and medium treatments was 100%. In contrast, birds in the high-dose group had a significant reduction in survival by 50% (χ² (3) = 14.94; p = 0.002, Fig. S1). Four individuals from the high dose died between days 3 and 12 of exposure, after ingesting an average daily dose (mean ± S.E.) of 704 ± 0.038 mg TMX/kg BW. No significant differences between sexes were observed in the survival of birds from high group (χ² (1) = 1.085; p = 0.298). The birds that survive were all males.

Thiamethoxam is classified as moderately to slightly hazardous (FAO, 2021). Acute toxicity information of TMX to birds is limited (Table S2). LD_50_ varies between 576 and 4366 mg TMX/kg BW, while dietary LD₅₀ exceeds 5200 mg TMX /kg bw, following an eight-day exposure through food. Although the current study was not designed to define acute dietary toxicity, the average dose that caused 50% of mortality in birds from the high treatment (i.e., 704 mg TMX/kg BW), can be used as a proxy due to the lack of data in passerines, the experimental similarities with the standardized protocol for the estimation of dietary toxicity (OECD, 1984), and the relative higher toxicity in comparison with the reported values. The survival pattern of baywings was similar to those observed after exposure to imidacloprid treated seeds in red legged partridges (*Alectoris rufa*) and grayish baywings (*Agelaioides* badius), where mortalities occurred between the third and fifteen days (Lopez-Antia et al., 2015, 2013; Poliserpi and Brodeur, 2023). The consistency in the survival curves is relevant in a worst-case scenario, as mortality would be expected after the third day of exposure to neonicotinoids.

### 3.2 Seed consumption, feeding behavior and body weight

Seed consumption was significantly influenced by treatment (p= 0.0021), week (p < 0.001), and their interaction (p = 0.0055) (Figure 1A). Birds exposed to higher doses significantly increased the consumption of seeds in comparison to the other experimental groups (p= 0.009). From the beginning of the exposure high TMX birds increased their consumption in 40% the first week, to 60% the last week in comparison with their pre-exposure consumption (p< 0.001 for all comparisons). Medium and low TMX birds increased their consumption gradually, compared with their pre-exposure levels. Medium TMX bird’s intake increased significantly in the last two weeks (p< 0.02) with a maximum increase of 16% at the end of exposure, while the intake of low treatment was significantly higher in the third week reaching a maximum of 13.4% (p= 0.005). A time effect was observed for all the birds in the first week with a decrease in consumption (p= 0.036). No differences were detected in the consumption of control birds (p= 0.26), and no sex effect was detected (F (1)= 0.228, p= 0.637).

**Figure 1.**
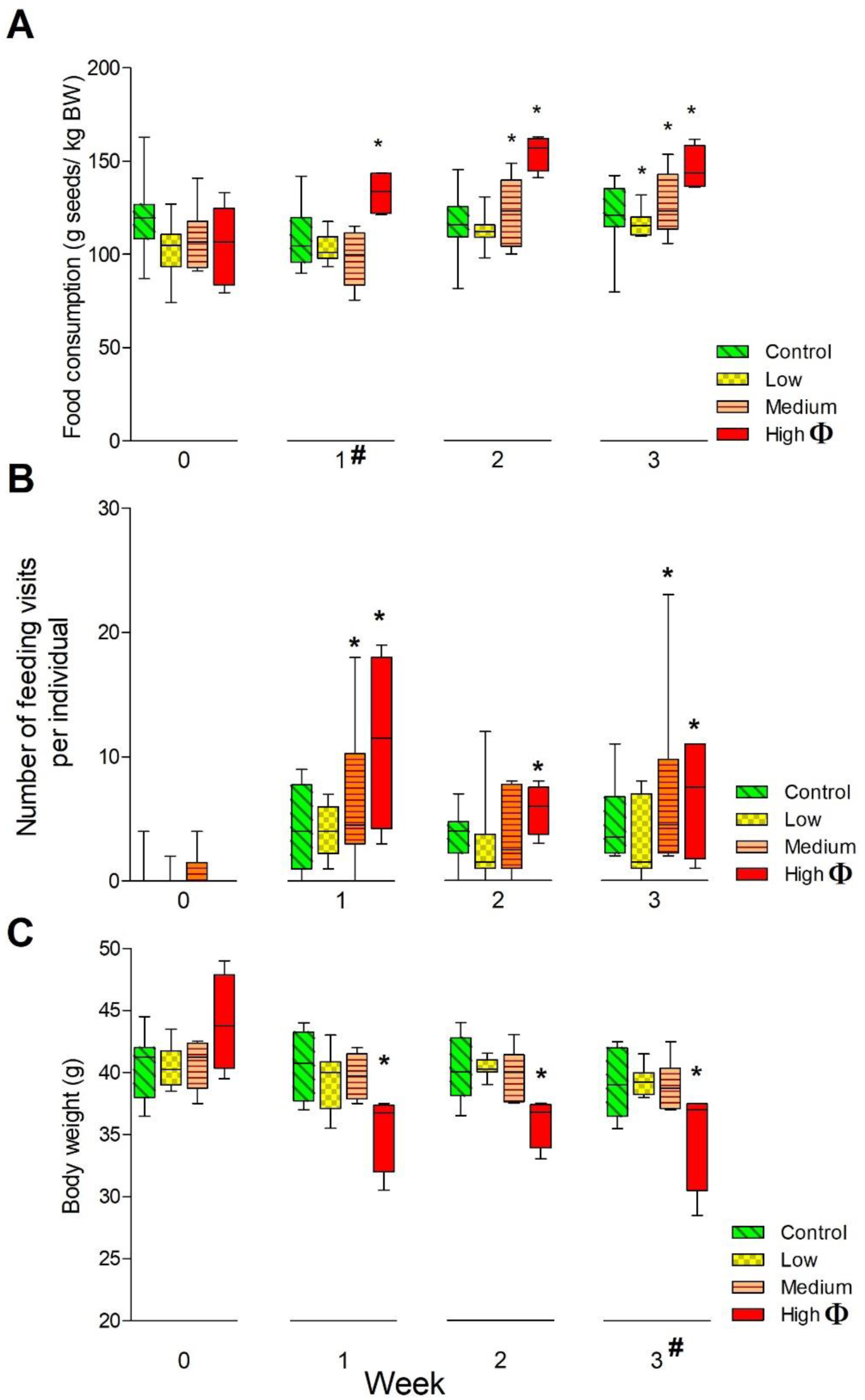
A) Weekly seed consumption, B) number of feeding events, C) body weight of *Agelaioides badius* exposed to TMX treated seeds. Significant differences (p<0.05): *= different from preexposure values; Φ= treatment effect; = week effect. Control: 0 g TMX/kg seed; Low: 0.027g TMX/kg seed; Medium= 0.334 g TMX/kg seed; High: 4.34 g TMX/kg seed. Boxes indicate interquartile range, middle lines indicate median, whiskers show the minimum and maximum values.

To our knowledge, this is the first study to report that birds exposed to TMX treated seeds increase their consumption in a dose-dependent way (Fig.S2: daily seed consumption). The literature on birds consuming neonicotinoids treated seeds is varied and may depend on multiple factors. In laboratory experiments, reported consumptions were equal or lower to controls depending on the dose: grayish baywings exposed to concentrations below 390 mg IMI /kg seed exhibit food intake levels like controls (Poliserpi et al., 2023); and the same was reported for red-winged blackbirds (*Agelaius phoeniceus*) and red-legged partridges (*Alectoris rufa*) exposed to 278 and 700 mg IMI/kg seeds, respectively (Avery et al., 1994; Lopez-Antia et al., 2013). Avoidance of treated seeds was reported for eared doves (*Zenaida auriculata*) exposed to 1.9 g TMX/kg or 4.5 g IMI/kg seed (Addy-Orduna et al., 2022), however the same author also reported the consumption of neonicotinoid treated-seeds by the same species in the wild (Addy-Orduna et al., 2025). Another influencing factor is that species may have different detoxification processes, and some could be more efficient than others at protecting themselves against toxic substances, therefore having different acceptance when exposed to neonicotinoids treated seeds (Arnold et al., 2015). Decrease in food consumption, or anorexia, caused by TMX and other neonicotinoids has been reported after exposure by gavage (Addy-Orduna et al., 2019; Eng and Morrissey, 2025; Shafia Tehseen Gul et al., 2020). The severity of these effects appears to depend on the exposure route. In baywings, the absence of anorexia can be explained by the fact that exposure occurred through food rather than gavage since the dose is ingested gradually and can be modulated by the birds’ behavior and physiology, resulting in a more continuous intake and absorption, unlike the rapid and concentrated dosing that occurs with gavage ((Poliserpi et al., 2021b, 2021a).

Figure 1B shows the weekly number of visits to the feeder. High TMX birds exhibited a general increase in the number of visits to the feeder (p= 0.007) in comparison with control group, and their pre-exposure values (p< 0.05 for all days). Compared with their pre-exposure values, medium TMX birds significantly increased the number of feeding events on weeks 1 and 3, (p< 0.01). In general, the number of feeding events was influenced by daytime (i.e., morning and afternoon) (p= 0.0266, Fig. S3). Comparison within treatments showed that control, low and medium birds had higher number of visits in the afternoon (p< 0.05), however birds from high TMX did not present differences in the number of feeding events regarding daytime (p= 0.65). Compared to control, high TMX birds also spend more time feeding (p= 0.0415, figure S4A), but no significant effect of daytime was observed within treatments (p= 0.18, figure S4B). No significant effects of treatment or daytime were observed in the duration of feeding events (p= 0.616, Fig. S5A and p=0.99, Fig. S5B; respectively).

Feeding behavior confirmed the lack of aversion and anorexia: birds exposed to TMX increased their number of feeding visits and their time feeding. In a similar experiment, Pan et al. (2022) reported that quails exposed to 20 and 200 mg TMX/kg seeds, displayed a positive sentiment for feeding although the authors did not quantify the amount of food ingested. In a previous study, we proposed that IMI neurotoxic effects may difficult the manipulation of food by increasing handling time (Poliserpi et al., 2023). However, no differences were detected in the duration of the feeding events in this case. Interestingly, the increase in the frequency of visits observed in birds from higher treatment had an effect in their diurnal feeding patterns. While all groups exhibited a significantly greater number of visits to the feeder in the afternoon, high-TMX exposed birds presented a constant feeding activity, with comparable consumption levels in both the morning and afternoon periods. The modification of the feeding pattern observed in baywings agrees with a previous study where white-crown-sparrows (*Zonotrichia leucophrys*) exposed to sublethal doses of IMI (1.2 or 3.9 mg/kg BW) experienced extended stopovers during migration, related to reduced fuel loads, suppression of feeding, and fueling ability (Eng et al., 2019). As was observed in baywings, the authors hypothesized that the longer stopovers were a result of birds needing more time to regain fuel stores through feeding. Regulation of appetite and food intake in birds represents a complex mechanism involving multiple levels of control that can be affected by neonicotinoid exposure (Costas-Ferreira and Faro, 2021). Because neonicotinoids are structurally similar to nicotine, they are expected to have similar effects on suppressing the appetite and food intake through agonism of nAChRs located in central and peripheral nervous system (Jo et al., 2002). However, neurotransmitters effects may depend on bird species and their physiological state: the effects can be opposite or have no effect (Denbow, 1999). Recently, Pan et al. (2024) informed abnormal changes in Japanese quails (*Coturnix japonica*) neurotransmitter levels related to food regulation, like dopamine and γ-aminobutyric acid (GABA), after the exposure to 21 and 210 mg TMX/kg seed for 28 days, which may support the impairment in feeding behavior and food intake observed in baywings.

The body weight of birds was significantly influenced by treatment (p= 0.009) and week (p = 0.016) (Figure 1C). Specifically, birds fed with the highest dose of TMX-treated seeds exhibited a significantly decrease in body mass, in comparison with the other experimental groups, and their initial weight (p < 0.001). Birds from high treatment lost an average of 20.5 % of their initial body weight, which remained decreased to the end of the experiment. The effect of the week was statistically significant in the third week, where a general decrease in body weight was observed for all the experimental groups (p = 0.035), but no significant changes in body weight were observed for control, low and medium treatments (p> 0.05). Body weight was not influenced by sex (p= 0.100). No statistically significant differences were observed in terms of the hepatosomatic index (HSI) when comparing among treatments and sex (F(3)= 0.826, p = 0.493 and F (1)= 1.455, p = 0.241; respectively).

The baywings exposed to TMX maintain their body weight or reduce it, contrary to what was expected based on consumption and feeding behavior. Previous studies have reported body weight loss, often linked to transient anorexia and increased energy expenditure due to detoxification processes (Eng and Morrissey, 2025; Lopez-Antia et al., 2015; Poliserpi and Brodeur, 2023). However, the change in body weight may result from a range of physiological mechanisms disrupted by TMX (Costas-Ferreira and Faro, 2021). Evidence suggests that neonicotinoids can disrupt thyroid function affecting basal metabolic rate and seasonal body weight homeostasis: lizards exposed to TMX and other neonicotinoids identified thyroid system disruption with distinct toxicity mechanisms (Wang et al., 2020), while red munias exposed to IMI exhibited thyroid histopathology and reduced hormone levels (Pandey & Mohanty, 2015). The impact of neonicotinoids on vertebrate body weight appears to be non-linear: in rats, the exposure to 0.5 or 1.0 mg IMI/kg BW for 60 days resulted in either weight gain or weight loss, respectively; likely due to pancreatic dysfunction and altered glucose metabolism (Khalil et al., 2017). Similarly, zebra finches (*Taeniopygia guttata*) exposed to 0.205 mg IMI/kg BW during early development, showed improved body condition in adulthood, characterized by increased lean mass and elevated basal metabolic rates (Zgirski et al., 2021). In baywings, BW appears to be less sensitive to TMX effects than food consumption or behavior, as was reported for red-winged blackbirds (Eng and Morrissey, 2025). The lack of weight variation, however, does not necessarily reflect good condition as some organ inflammatory response may be occurring (Elhamalawy et al., 2022; Ma et al., 2022; Tokumoto et al., 2013). Although no differences in HSI were detected, other organs may be affected.

### 3.3 Anti-predatory behavior, jumping and cholinesterase activity

TMX significantly affected the anti-predatory behavior in birds exposed to the highest doses of TMX, that exhibited a general decrease in the reaction (p=0.0067) and intensity to the stimulus (Fig. 2, p<0.006). No significant differences were observed in anti-predatory behavior of low and medium groups, although both groups reduced their reaction by 11% compared to controls. No significant effects of week or the interaction week*treatment were found on the reaction and intensity (p> 0.05).

**Figure 2.**
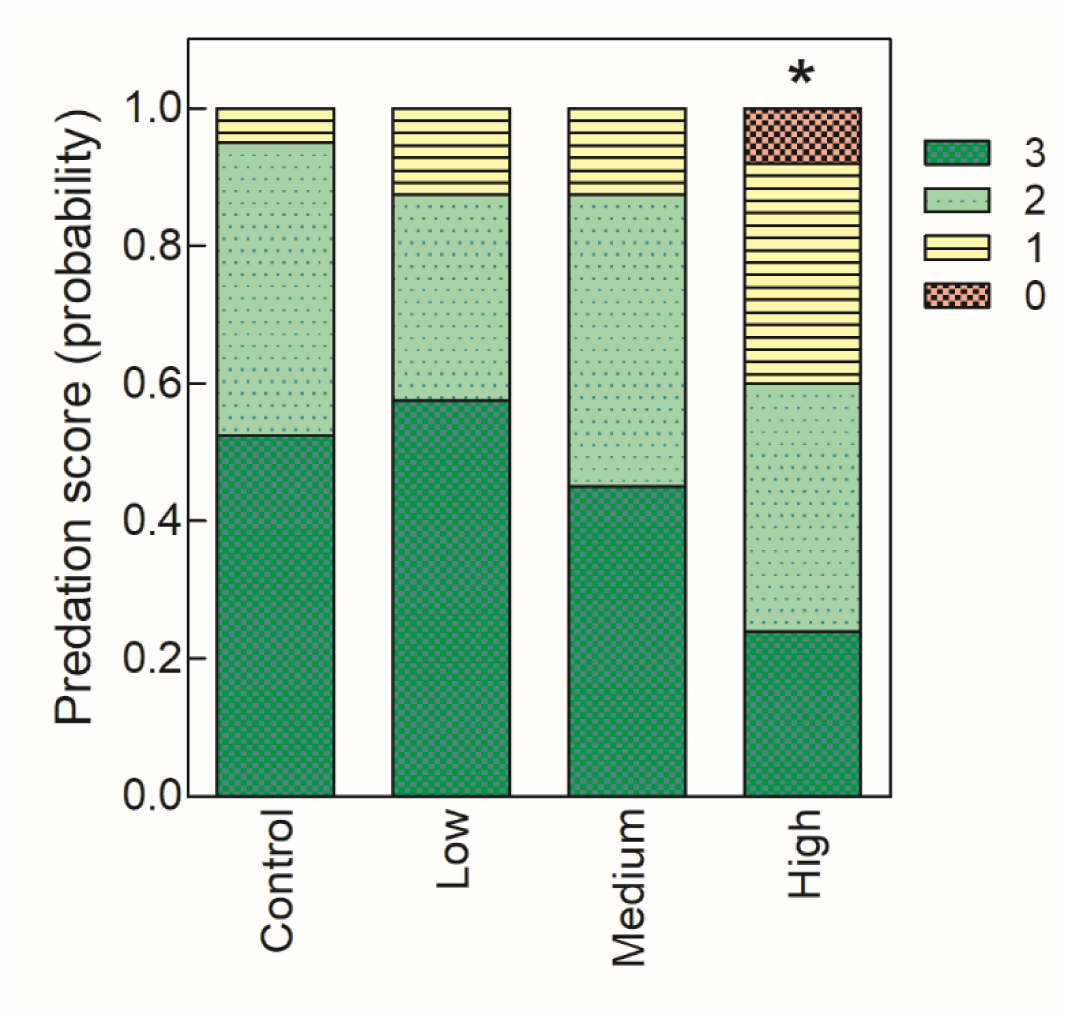
Intensity of the anti-predatory response of *Agelaioides badius* exposed to TMX treated seeds, where 0=none and 3=very strong. *= significantly different from control (p < 0.05). Control: 0 g TMX/kg seed; Low: 0.027g TMX/kg seed; Medium=0.334 g TMX/kg seed; High: 4.34 g TMX/kg seed.

The effect of neonicotinoids on the impairment of the anti-predatory behavior was previously reported in birds (Moreau et al., 2022) and other vertebrates (Lee-Jenkins and Robinson, 2018) (Faria et al., 2020). Single exposure to IMI (i.e.,6 mg/kg bw), caused red legged partridges to exert hyperreactivity (i.e., increased alert behavior), but difficulty in staying crouched when presented with predator decoys (Addy-Orduna et al., 2024); while continuous exposure to IMI (i.e., 514 mg IMI/kg BW) reduced the response of grayish baywings to a predator vocalization and to an approaching person (Poliserpi and Brodeur, 2023). The ability of birds to respond to ecological stress is mediated in large part by glucocorticoids involved in metabolic, cardiovascular, immunologic and homeostatic functions (Burger et al., 2015; Crespi et al., 2013). Under predation risk, there is an increase in the production of glucocorticoids to mobilize energy and resolve stressful stimuli by modulating the intensity of behavioral responses (Fontaine et al., 2011; Scheuerlein et al., 2001; Voellmy et al., 2014), that may be disrupted by neonicotinoids (Gavel et al., 2019; Wang et al., 2018). Ecologically, predator avoidance plays a critical role in key behavioral decisions like exploration (Huang et al., 2012) and nest site selection (Enos et al., 2023) that may be adversely affected by exposure to TMX.

All the baywings showed a higher number of jumps in the morning compared to the afternoon, regardless of the treatment (*p*< 0.001, Figure S6). Figure 3 illustrates the total number of jumps per treatment over the exposure period. There was a general effect of time (p< 0.001) where all treatments decreased their frequency of jumps at the end of the experiment, and a significant interaction between treatment and day (p< 0.001), so comparisons were made within each treatment versus their preexposure period. Birds from high treatment experienced a general reduction in the number of jumps (*p*= 0.0293), all days being significantly lower than initial values (p<0.05). Medium treatment exhibited a similar tendency (*p*= 0.0794), and the decrease was significantly differente the firs week (p < 0.001). Birds from the low treatment, however, increased the number of jumps on the second and third weeks (*p*< 0.001). No statistically significant differences were observed within the control group (p> 0.05).

**Figure 3.**
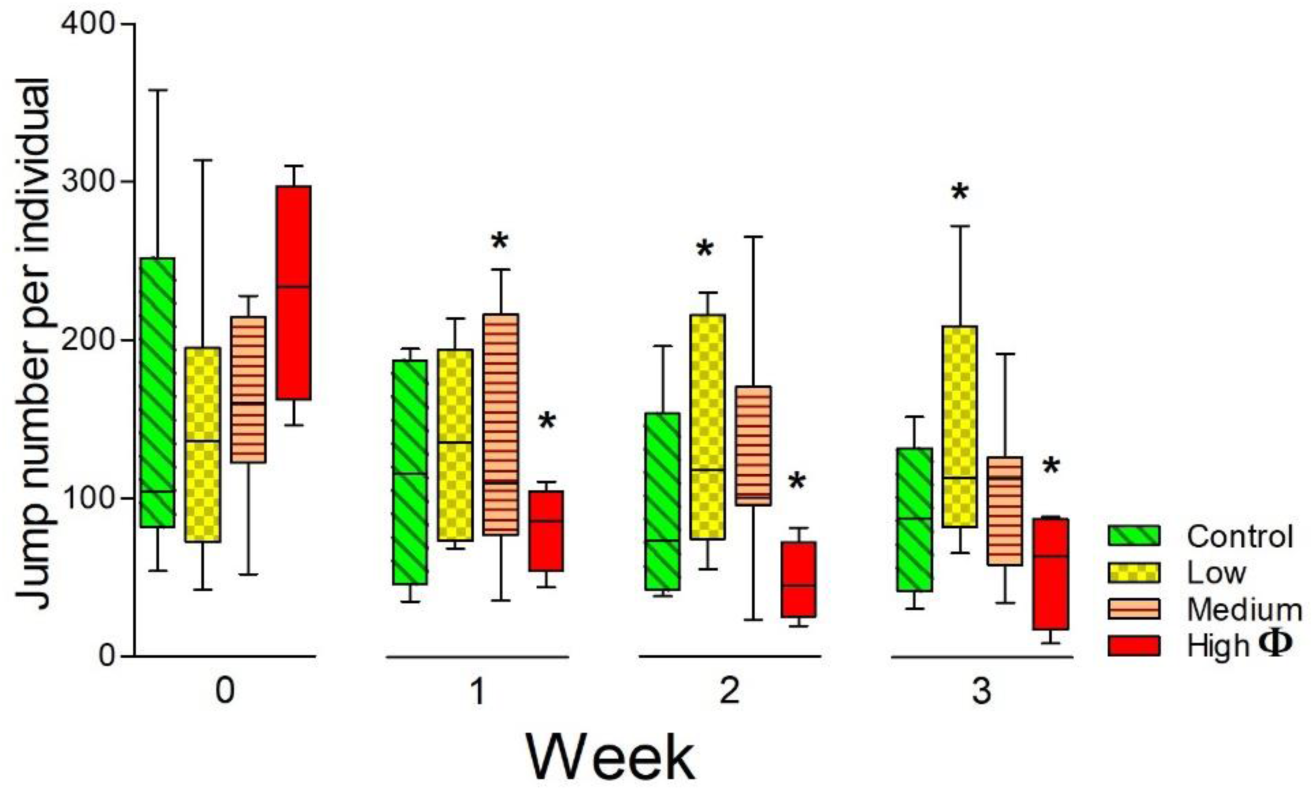
Number of jumps (Mean ± S.E.) of *Agelaioides badius* exposed to TMX treated seeds. *= significantly different from preexposure values within treatment; Φ= treatment effect (p < 0.05). Control: 0 g TMX/kg seed; Low: 0.027g TMX/kg seed; Medium= 0.334 g TMX/kg seed; High: 4.34 g TMX/kg seed. Boxes indicate interquartile range, middle lines indicate median, whiskers show the minimum and maximum values.

Jump frequency can be considered an indicator of locomotor activity. Notably, baywings exposed to lower doses of TMX increased their activity at the end of the exposure, contrary to the general decrease exhibited by birds from medium and high treatments. Impairment of locomotor activity or mobility due to neonicotinoids exposure was related to its neurotoxic effect in vertebrates (Costas-Ferreira and Faro, 2021; Gibbons et al., 2015). Hyperactivity due to TMX was related to anxiety-like behavior: in rats defined as the decrease in the proportion of time and entries in open spaces in a maze (Rodrigues et al., 2010), while zebra fishes increased their average speed (Zhang et al., 2021). Hypoactivity, on the other hand, was extensively reported in birds, and is often related to loss of balance, uncoordinated movement, and quiescence (Eng et al., 2019; Franzen-Klein et al., 2020; Poliserpi et al., 2021a). The effects of neonicotinoids on sensorimotor functions in vertebrates were related to alterations in neuronal activity and neurotransmitters that regulates the circuitry underlying locomotor behavior (Sholomenko et al., 1991; Wang et al., 2018). In Japanese quails, 4.5 mg IMI/kg BW induced a significant decrease in dopamine and significantly elevated norepinephrine and serotonin levels, resulting in disorders of cognition and behavioral changes characterized by excitability (Rawi et al., 2019). On the contrary, rats exposed to 1 mg IMI/kg BW for 60 days, exhibited less exploratory activity, deficit in sensorimotor functions, and depression; these effects associated with reduced levels of serotonin, gamma-aminobutyric acid, and dopamine (Abd-Elhakim et al., 2018).

Figure 4A shows the enzymatic activity of ChE measured in the brain, muscle, liver, kidney and plasma. Che activity was generally increased depending on the treatment and sex: in brain Che activity was elevated in low treatment (p = 0.0379), while in the liver, only males from medium treatment showed significant differences (p=0.016). In muscle birds from high (p= 0.002) and females of low (p= 0.041) treatments showed an increase in enzymatic activity. Only birds from high treatment had an increase in Che activity in plasma (p=0.043). ChE activity did not present any further significant differences among treatments or sex in kidney (p>0.05).

**Figure 4.**
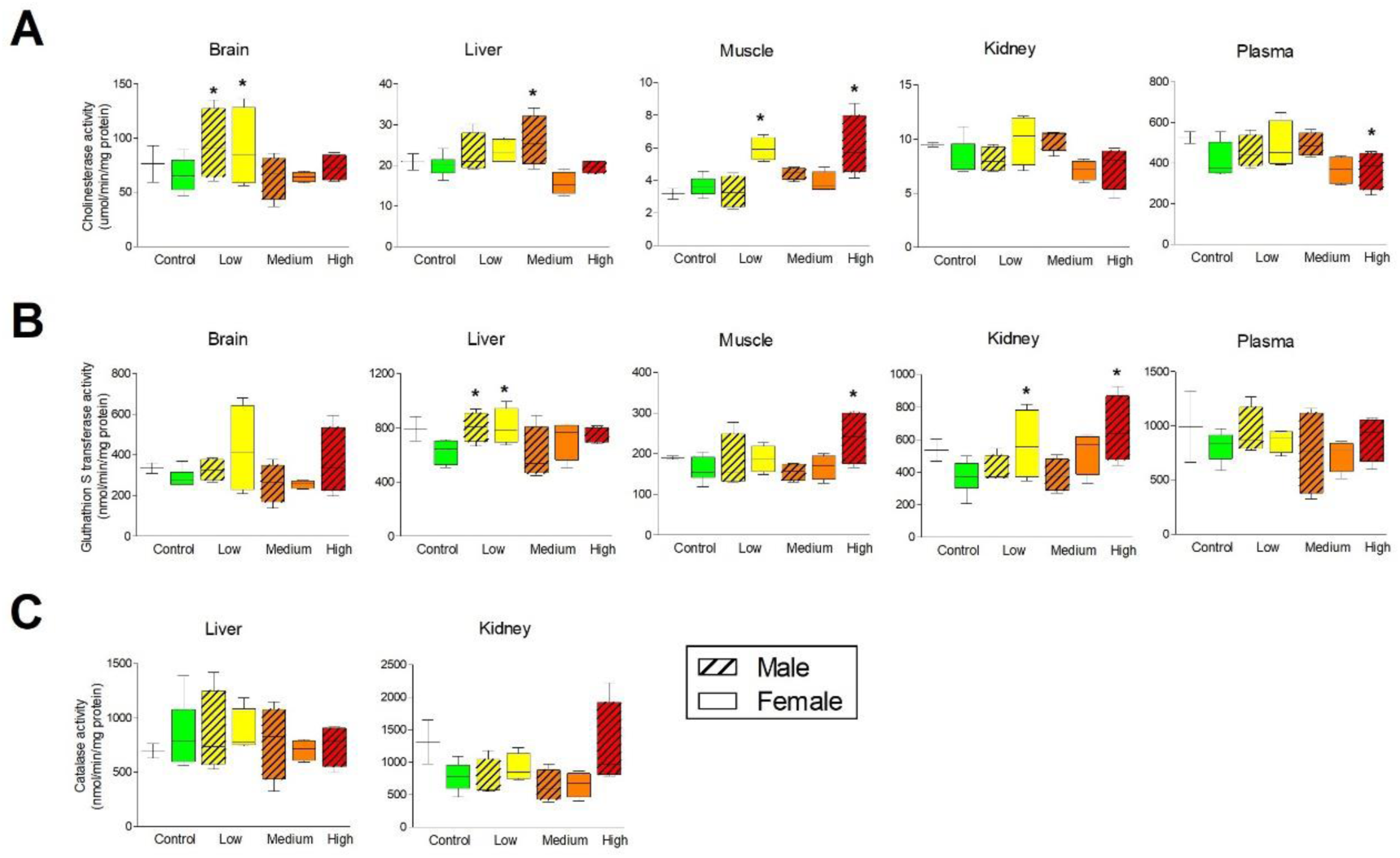
Enzymatic activity (Mean ± S.E.) of A) Cholinesterases, B) gluthathion S transferase and C) Catalase of *Agelaioides badius* exposed to TMX treated seeds. *= significantly different (p < 0.05). Control: 0 g TMX/kg seed; Low: 0.027g TMX/kg seed; Medium= 0.334 g TMX/kg seed; High: 4.34 g TMX/kg seed. Boxes indicate interquartile range, middle lines indicate median, whiskers show the minimum and maximum values.

The cholinergic system plays an important role in locomotor activity, particularly, the acetyl-cholinesterase (AChE) is responsible for breaking down acetylcholine at both the neuromuscular junction and within the central nervous system (Scanes, 2014). Dysregulation of AChE may lead to the accumulation of acetylcholine (Ach), resulting in neuromuscular dysfunction and tremors (McMahan et al., 1978; Nassar, 2016). The general increase of ChE activity (i.e., which include AChE activity) of exposed baywings, agrees with reported effects in vertebrates that propose that ChE activity may increase or decrease as a compensatory mechanism (Wang et al., 2018; Zhang et al., 2021). It was recently proposed for lizards (*Eremias argus*) that TMX do not work on nAChRs but directly increase the concentrations of ACh by up-regulating the expression of the ACh gene, which in turn enhances the binding of ACh and nAChRs (Wang et al., 2019). Because AChE is involved in terminating the nervous impulse it is possible that a regulation pathway can be activated, after TMX exposure (Schweitzer, 1993). Although a sex effect was detected in the enzymatic activity, this was probably due to a sample size as no clear pattern was detected, and no sex effect were previously reported in baywings ecotoxicity tests (Fernández-Vizcaíno et al., 2025; Poliserpi et al., 2021a).

### 3.4 Enzymatic activity and hematology

The enzymatic activity of GST is shown in Figure 4B. GST was increased in liver, muscle and kidney depending on treatment and sex. In the liver, only birds from low treatment had a general increase in GST activity (p=0.0356), while in the muscle, activity was increased in birds from high treatment (p= 0.0177). In the kidney GST activity was increased in birds from high treatment (p=0.013), in females from low group (p=0.0492) and the same tendency was observed in females of medium group (p=0.0588). No statistically significant differences were detected in GST activity in brain and plasma (p>0.05); or in CAT activity (p> 0.05, Fig. 4C).

GSTs are a family of phase II detoxification enzymes that play a central role in the biotransformation of xenobiotics and the maintenance of cellular redox homeostasis (Newman, 2015). In general, oxidative stress is among the main ways by which neonicotinoids cause tissue damage and the evidence points to the role of GST as part of antioxidant system (Pan et al., 2022a, 2022b; Wang et al., 2018) therefore, the general increase in GST activity observed may be indicative of an imbalance in baywing’s redox system. Pan et al. (2022a) evidenced, in a similar experiment, that transcriptomic and metabolomic analyses of TMX exposed quails caused changes in Glutathione (GSH) metabolism, a non-enzymatic substrate of GST, that ultimately may impair GST response to oxidants. Moreover, the authors proposed different TMX mechanisms of oxidative stress, depending on the dose: low dose (i.e. 20 mg/kg BW) caused oxidative DNA damage and high dose (i.e., 200 mg/kg BW) resulted in ferroptosis, a regulated necrotic cell death, and suppression of antioxidant capacity (Pan et al., 2022a). Oxidative stress was also evidenced by Gul et al., (2020) that exposed broiler chicks to 100 mg TMX/kg BW combined with the antioxidant selenium and vitamin E and observed a general amelioration of the effects. Supporting these results, administration of antioxidants improved the effects exerted by mice dosed with 87.73 mg TMX/kg BW (Elhamalawy et al., 2022). Although no effects were detected in CAT activity, it must be taken into account that the antioxidant capacity varies among species, tissues and cell types (Costantini, 2008).

Hematological parameters determined in baywings at the end of the exposure are presented in figure 5. Significant dose-dependent effects were found for PCV (p < 0.001; Fig.5A), while polychromasia levels were only decreased in high TMX birds (p = 0.041, Fig.5B). No significant effect of treatment was detected in WBC (p>0.05), and no sex effect was observed in any hematological parameters (p>0.05).

**Figure 5.**
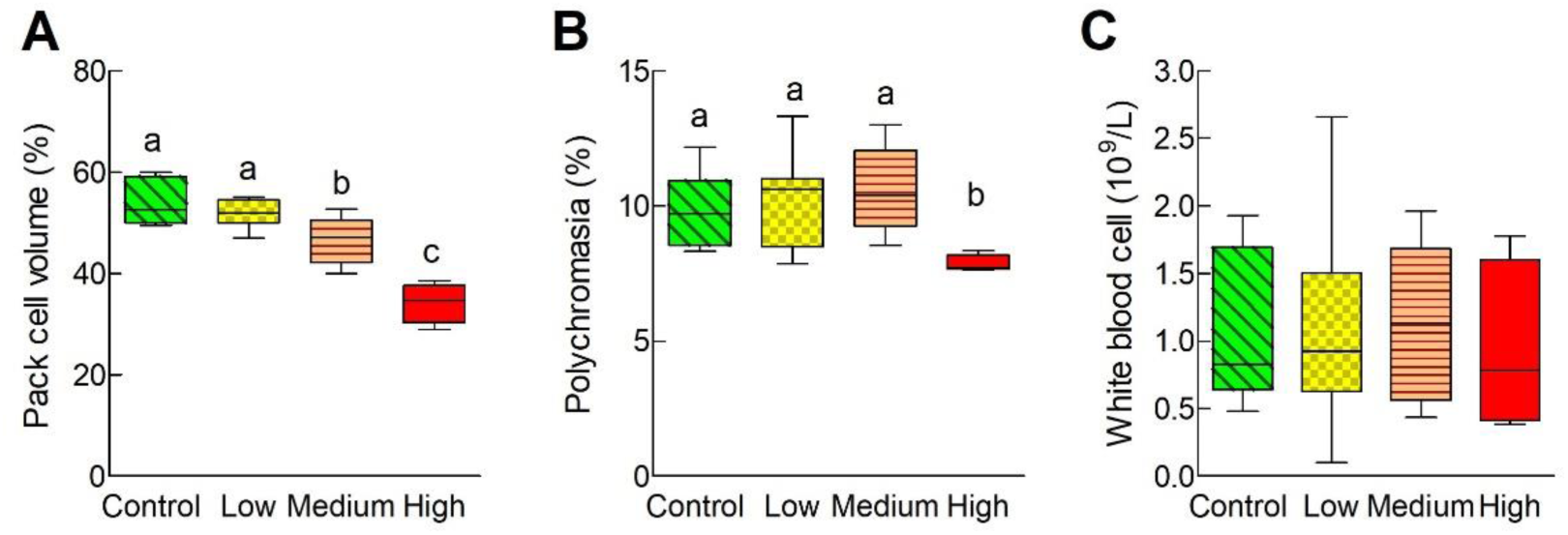
Hematological parameters (mean ± S.E.) of *Agelaioides badius* after exposure to TMX: A) Pack cell volume, B) Polychromasia and C) White blood cell count after exposure to TMX treated seeds. Different letters indicate significant differences (p<0.05). Control: 0 g TMX/kg seed; Low: 0.027g TMX/kg seed; Medium= 0.334 g TMX/kg seed; High: 4.34 g TMX/kg seed. Boxes indicate interquartile range, middle lines indicate median, whiskers show the minimum and maximum values.

A dose-dependent anemia was observed, indicating a decreased number of circulating erythrocytes. TMX and its main metabolite CLO were reported to alter hematological profiles in vertebrates (Elhamalawy et al., 2022; Yaseen et al., 2025). Gul et al, exposed either boiler chicks or laying hens to doses between 50 and 1000 mg TMX/kg BW for 45 days, that resulted in a reduction in erythrocyte count, hemoglobin and hematocrit (Gul et al., 2020; Gul et al., 2019, 2018), and associated the abnormal hematological profile to oxidative damage (Gul et al., 2020). Nonregenerative anemia occurs when new red blood cells are not produced and released into circulation: wood frogs (*Lithobates sylvaticus*) chronically exposed to clothianidin (2.5 µg/L) and thiamethoxam (2.5 or 250 µg/L) remained anemic 3 weeks after exposure to neonicotinoids (Gavel et al., 2019). In this study polychromasia was assessed as a measure of blood cell proliferation (Clark et al., 2009). The joint decrease in PCV and polychromasia in baywings exposed to high dose of TMX suggests non regenerative anemia. This might be due to the effect of TMX on the hemopoietic system: in birds erythropoiesis occurs in bone marrow and the proliferation of avian erythroblasts is promoted by erythropoietin in the kidney (Scanes, 2014) Previous studies in birds have not explored these hematological mechanisms specifically, but evidence points to TMX induction of oxidative stress causing abnormalities in bone marrow and kidney and the disruption of erythropoietin functions (Gavel et al., 2019; Wang et al., 2018). No differences were detected in WBC values after exposure to TMX, in contrast with previous studies where broiler chicks exposed to TMX decreased their WBC count after 42 days ( Gul et al., 2020, 2018). These results are consistent with previous reports that highlight that species-specific responses to contaminants have become more apparent, particularly regarding the hematological profile (Gavel et al., 2021, 2019; Poliserpi and Brodeur, 2023).

## CONCLUSION

This study provides evidence that repeated ingestion of TMX-treated seeds, at environmentally-relevant doses, can trigger significant behavioral, physiological, hematological, and biochemical alterations in grayish baywings. In a scenario where birds gorge on spilled seeds, mortality can occur as was observed in birds exposed to the highest dose. A dose-dependent increase in seed consumption was observed in the latter stages of the experiment, which amplifies their exposure risk. Small birds like grayish baywings (< 100g) have metabolic demands inherently higher compared with larger species, therefore exhibit proportionally greater food requirements and lower energy reserves (Muller et al., 2024). Altogether the increase in food consumption and foraging behavior, with the loss or non-gain of BW, suggest that TMX exposure may reduce energy reserves affecting the survival of birds in the wild (Cooper et al., 2015; Klaassen et al., 2004; Laursen et al., 2019). Bird’s survival also depends on the locomotor activity and their capacity to avoid predators, which were impaired by TMX (Muller et al., 2024; Saaristo et al., 2018). At the biochemical level, the alteration in hematological profiles probably related to oxidative stress can be detrimental because the reduction in oxygen and carbon dioxide–carrying capacities can result in tissue damage, decreased fitness, and increased stress (Clark et al., 2009; Scanes, 2014). Given that tested doses can be easily reached by birds by consuming less than 10% of their daily diet as treated seeds, these findings underscore the need to reconsider these effects in current risk assessment frameworks.

## Supporting information

Concentrations of TMX used for seed coating in Argentina, reported TMX LD50 for birds, additional figures of survival, seed ingestion, feeding behavio

## SUPPORTING INFORMATION

Concentrations of TMX used for seed coating in Argentina, reported TMX LD_50_ for birds, additional figures of survival, seed ingestion, feeding behavior and activity (i.e. jump).

## ACKNOWLEDGEMENTS

The current study was funded by the “Instituto Nacional de Tecnología Agropecuaria” (2019-PE-E2-I054-001). E. Fernández-Vizcaíno was financed by the Banco Santander Ibero-America Research Scholarships.

