## Supplementary material for "“Eating to Survive… or not? feeding compensation and toxicity in a songbird exposed to thiamethoxam treated seeds”": Concentrations of TMX used for seed coating in Argentina, reported TMX LD50 for birds, additional figures of survival, seed ingestion, feeding behavio

**Table S1.** Range and average of application rates of thiamethoxam (TMX) approved for seed treatment of common crops of the Pampa Region of Argentina.

|  | Application average  (g TMX/kg seed) | Application range^1^  (g TMX/kg seed) |
| --- | --- | --- |
| *Alfalfa* | 1.80 | 1.80 |
| *Oat* | 1.10 | 0.70 - 1.50 |
| *Sunflower* | 1.99 | 1.87- 2.10 |
| *Corn* | 2.10 | 1.20 - 3.00 |
| *Soybean* | 0.54 | 0.35 - 0.72 |
| *Sorghum* | 1.72 | 1.50 - 1.93 |
| *Wheat* | 0.36 | 0.36 |
| *Pasture* | 1.10 | 0.70 - 1.50 |
| **Total** | **1.37** | **0. 36 – 3** |

^1^.(SENASA, 2024)

SENASA, 2024. Registro nacional de terapéuticaVegetal: Vademecum [WWW Document]. URL https://aps2.senasa.gov.ar/vademecum/app/publico/formulados (accessed 11.5.25).

**Table 2.** Acute toxicity (LD50 and LC50) of thiamethoxam (TMX) in birds. Studies are grouped according to whether the experiments were performed with technical TMX or formulated.

| **Common Name** | **Latin Name** | **Exposure** | **Tested TMX** | **Endpoint** | **Concentration** | **Expected NOAEL 1/10 th** | **Days** | **Reference** |
| --- | --- | --- | --- | --- | --- | --- | --- | --- |
| Mallard duck | *Anas platyrynchos* | Food | Technical  98,6% | LC50 | > 5200  ppm | 520 | 8 | Environmental Fate and Effects Division, U.S.EPA, Washington, D.C. (1992) |
| Northern bobwhite quail | *Colinus virginianus* | Food | Technical  98,6% | LC50 | > 5200  ppm | 520 | 8 | Environmental Fate and Effects Division, U.S.EPA, Washington, D.C. (1992) |
| Mallard duck | *Anas platyrynchos* | Oral via capsule | Technical  98,6% | LD50 | 576 mg/kg bw | 56.7 | 14 | Environmental Fate and Effects Division, U.S.EPA, Washington, D.C. (1992) |
| Northern bobwhite quail | *Colinus virginianus* | Oral via capsule | Technical  98,6% | LD50 | 1552 mg/kg bw | 155 | 21 | Environmental Fate and Effects Division, U.S.EPA, Washington, D.C. (1992) |
| Eared dove | *Zenaida auriculata* | Gavage | Cruiser 60 FS* 60% | LD50 | 4366 mg/kg bw | 437 | 3 | Addy-Orduna *et al*., 2019 |

*Flowable concentrate for seed treatment.

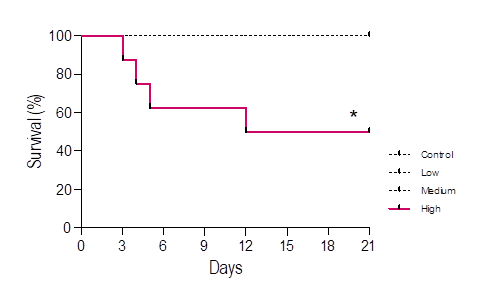

**Figure S1**. Survival of *Agelaioides badius* exposed to TMX treated seeds over 21 days. *= significantly different from control (p< 0.05). Control: 0 g TMX/kg seed; Low: 0.027g TMX/kg seed; Medium= 0.334 g TMX/kg seed; High: 4.34 g TMX/kg seed.

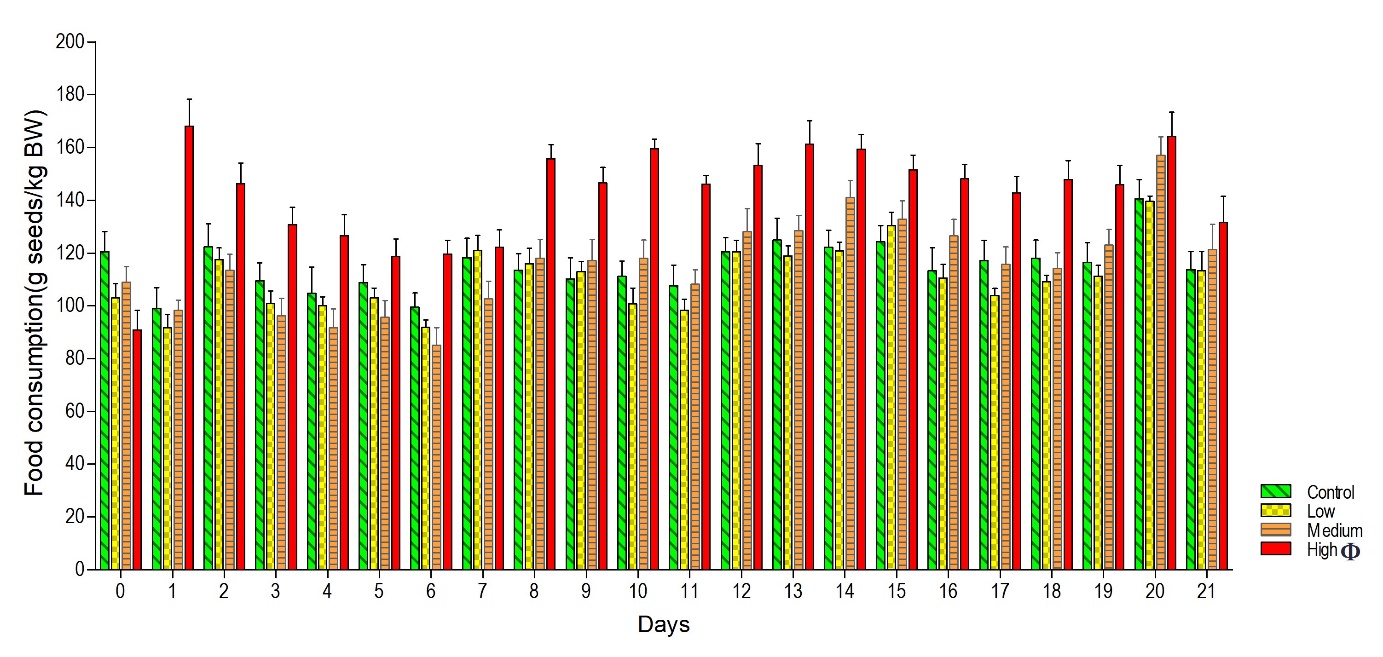
**Figure S2.** Daily seed consumption (mean ±SE) of *Agelaioides badius* exposed to TMX treated seeds. Φ= treatment effect (p < 0.05). Control: 0 g TMX/kg seed; Low: 0.027g TMX/kg seed; Medium= 0.334 g TMX/kg seed; High: 4.34 g TMX/kg seed.

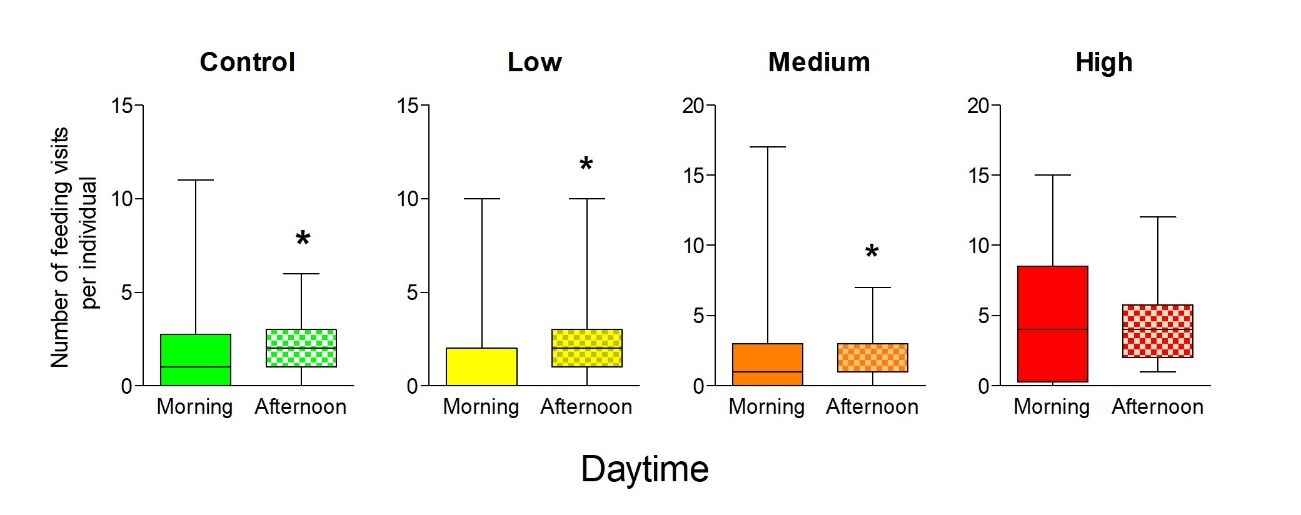
**Figure S3:** Daytime number of feeding visits (Mean ± S.E.) of *Agelaioides badius* exposed to TMX treated seeds. *= significantly different (p < 0.05). Control: 0 g TMX/kg seed; Low: 0.027g TMX/kg seed; Medium= 0.334 g TMX/kg seed; High: 4.34 g TMX/kg seed.

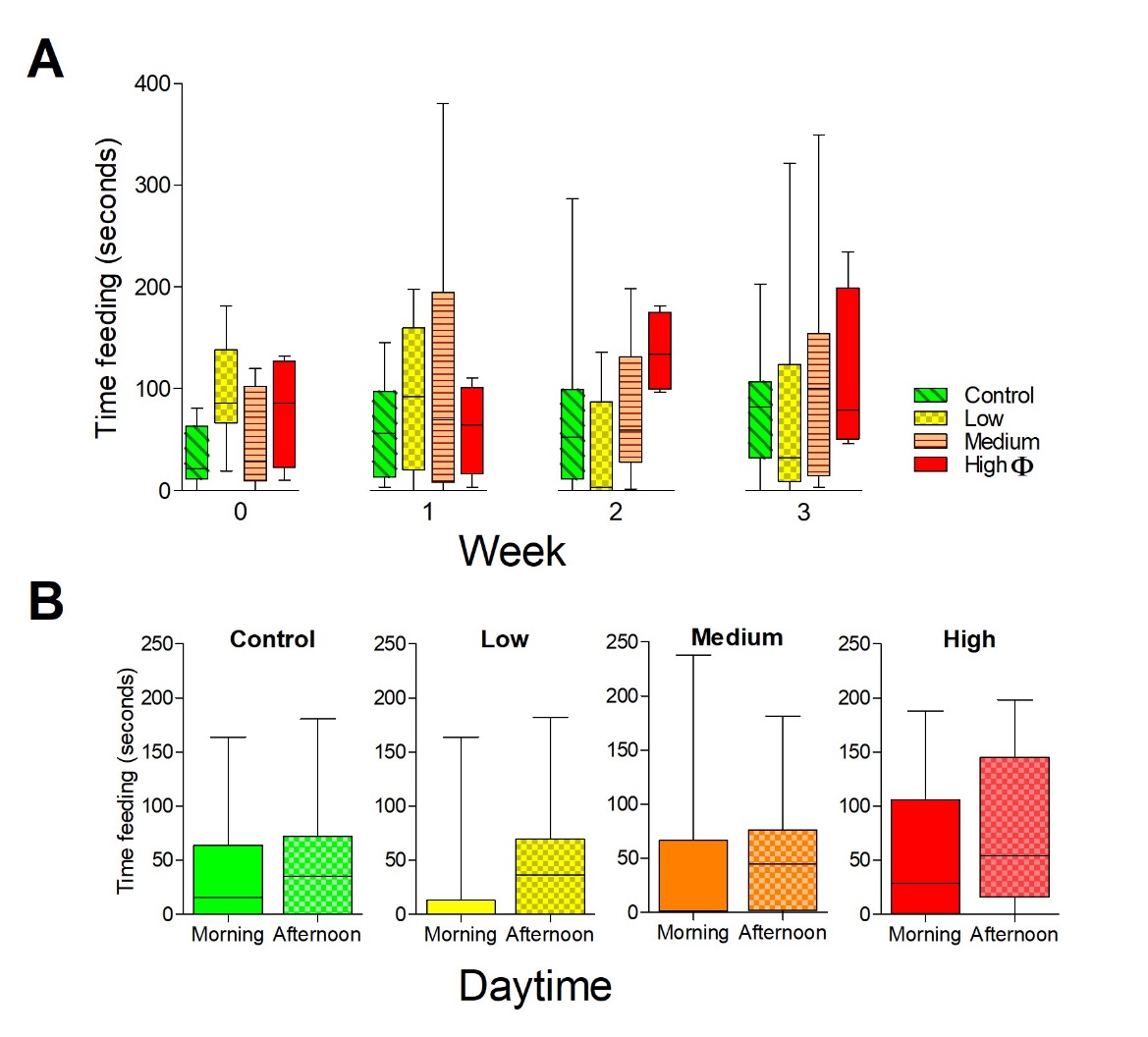

**Figure S4:** A) total time feeding and B) time feeding within treatments according to daytime of *Agelaioides badius* exposed to TMX treated seeds. Φ= treatment effect (p < 0.05) Control: 0 g TMX/kg seed; Low: 0.027g TMX/kg seed; Medium= 0.334 g TMX/kg seed; High: 4.34 g TMX/kg seed. Boxes indicate interquartile range, middle lines indicate median, whiskers show the minimum and maximum values.

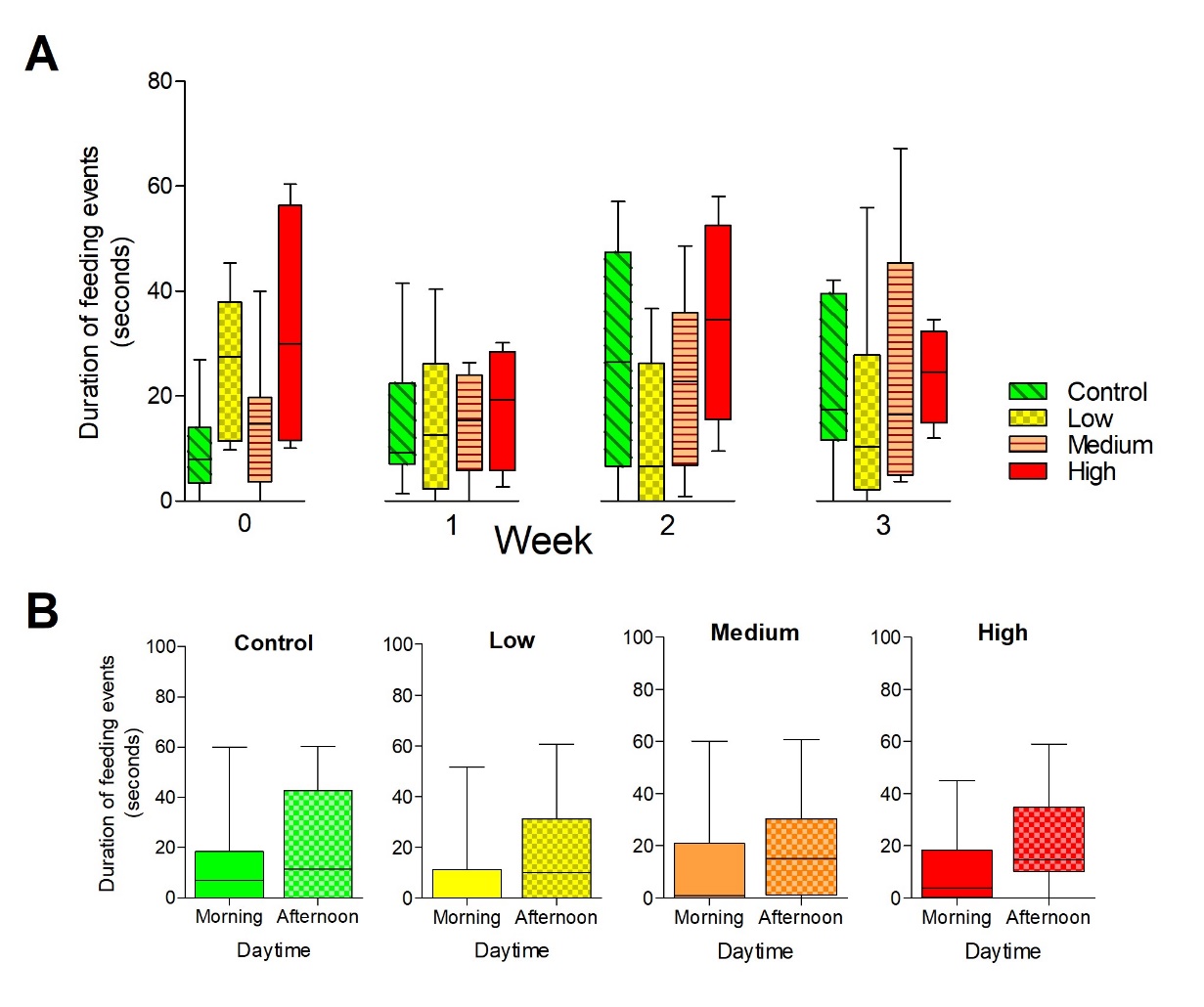
**Figure S5:** A) Mean duration of feeding events and B) Duration of feeding events within treatments according to daytime of *Agelaioides badius* exposed to TMX treated seeds. Control: 0 g TMX/kg seed; Low: 0.027g TMX/kg seed; Medium= 0.334 g TMX/kg seed; High: 4.34 g TMX/kg seed. Boxes indicate interquartile range, middle lines indicate median, whiskers show the minimum and maximum values.

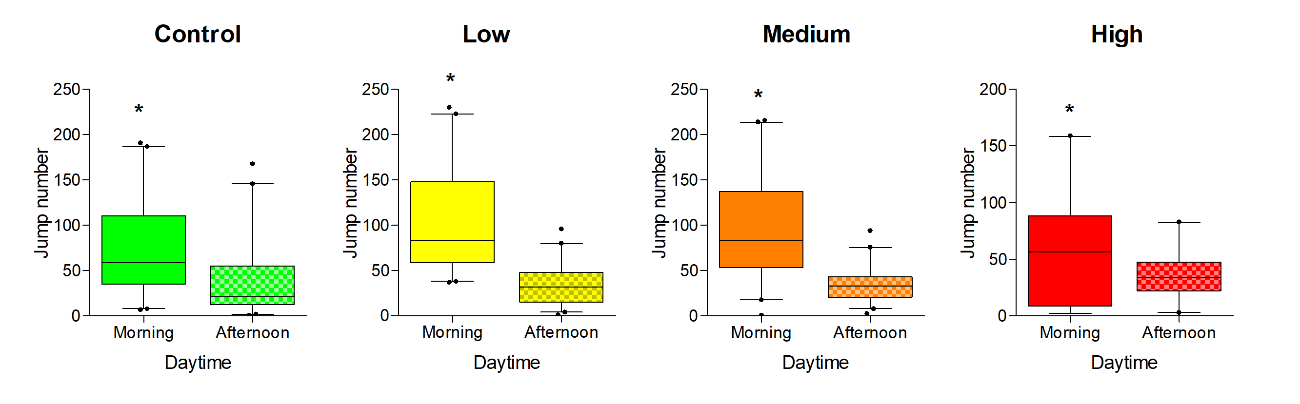
**Figure S6:** Daytime number of jumps of *Agelaioides badius* exposed to TMX treated seeds. *= significantly different (p < 0.05). Control: 0 g TMX/kg seed; Low: 0.027g TMX/kg seed; Medium= 0.334 g TMX/kg seed; High: 4.34 g TMX/kg seed. Boxes indicate interquartile range, middle lines indicate median, whiskers show the minimum and maximum values.
